# A validated method for banking mixed glial cultures that yield responsive microglia

**DOI:** 10.64898/2025.12.13.693524

**Authors:** Annalise M. Sturno, James E. Hassell, Katelyn M. Baker, Garrett J. Potter, Kimberley D. Bruce

## Abstract

Microglia, the primary immune cells of the brain, orchestrate immune responses to both external and internal stimuli in health and disease. Although several cell culture methods exist to model microglia *in vitro*, isolating loosely adherent microglia from primary murine mixed glial cultures remains valuable for recapitulating *in vivo* cell states from complex transgenic lines at relatively low cost. However, these methods are constrained by limited temporal control and scalability. To address these challenges, we established a protocol for generating microglia from frozen mixed glial cultures. While freezing introduced measurable differences in microglial morphology and lipid droplet content, microglia derived from frozen cultures responded to environmental stimuli (high glucose and LPS) in the same way as those from fresh cultures. Thus, while frozen primary microglia exhibit similar functional responses, maintaining consistency in using fresh or frozen cells within a given experiment is strongly recommended.

**Motivation:** Although our understanding of microglia’s role in brain development and disease continues to grow, a standardized *in vitro* model of primary mouse microglia remains absent in the field. Primary cell culture systems are essential for studying microglia, enabling the empirical assessment of gene function in transgenic models and testing novel interventions before committing to time- and cost-intensive *in vivo* studies. However, mixed glial culture-derived primary microglia (pMG) systems are constrained by the timing of animal availability and have limited scalability. In this study, we sought to establish a protocol for freezing and subsequently reviving primary mixed glial cultures from which responsive pMG could be isolated for downstream analysis. We validated this approach by quantifying microglia responses to distinct environmental stimuli before and after a freeze-thaw cycle.

**Research topics:** CP: Cell biology

**Graphical Abstract:** 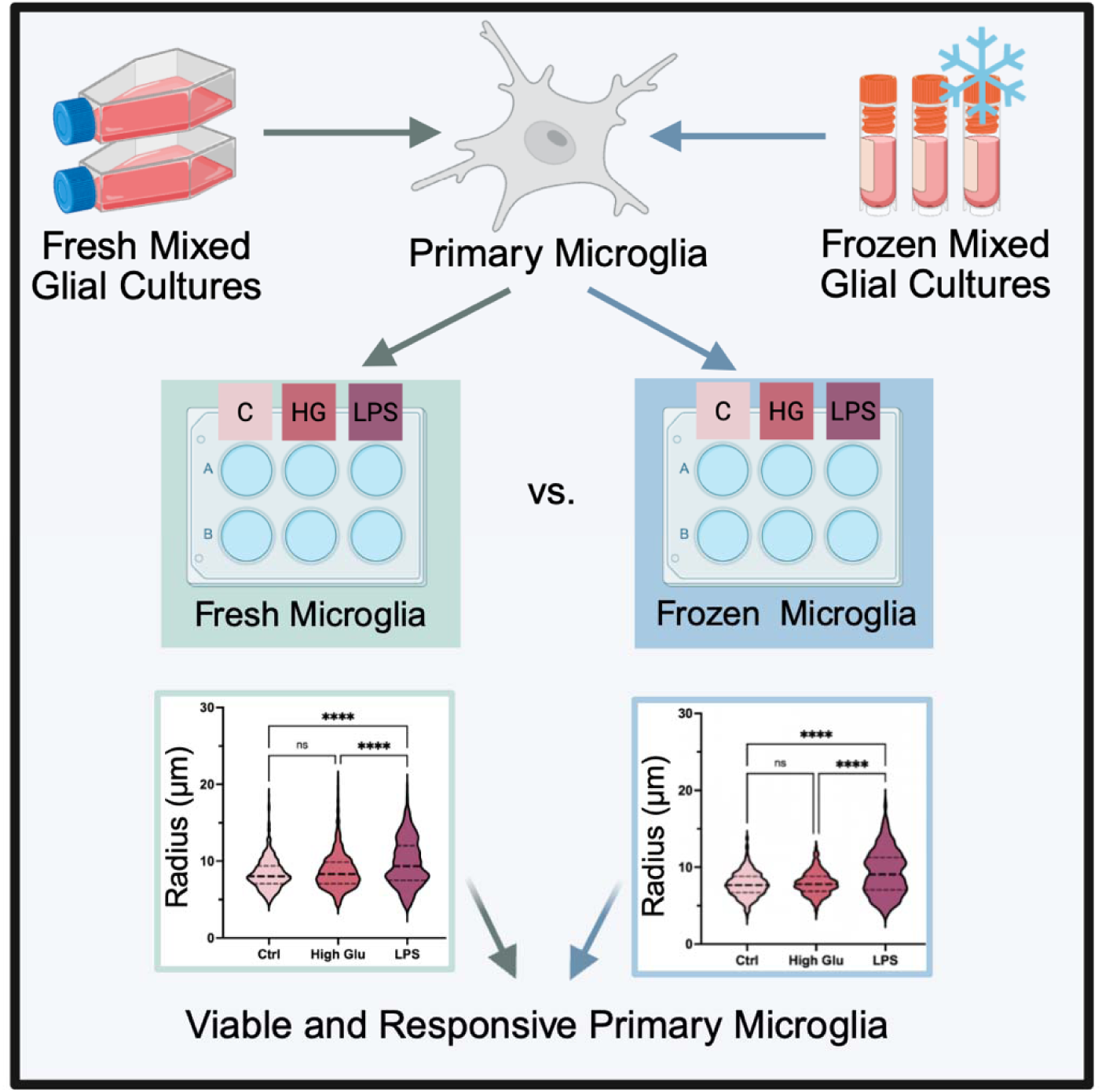

## Introduction

Microglia, the macrophages of the brain, are critical mediators of brain health and disease^1, 2^. Microglia regulate the brain’s immune system by rapidly responding to changing conditions and noxious stimuli. At their baseline, homeostatic state, microglia are highly motile and survey the brain in search of debris or inflammatory signals^1^. When encountering external stimuli, microglia rapidly respond, often becoming activated. Microglial activation has been identified as an essential process in directing natural aging^3^, responses to stress^4^, and combating disorders like Alzheimer’s disease^5^ and Parkinson’s disease^6^. While microglial activation exists on a spectrum of phenotypes, it is generally characterized by altered morphology^7^, increased phagocytosis^8^, and pro-inflammatory cytokine release^4^. However, with an increasing appreciation of microglial heterogeneity, changes in microglia state can also characterized by their metabolism^5^. Specifically, metabolic changes in activated microglia be are reminiscent of the Warburg effect, where metabolism is shifted away from oxidative phosphorylation towards glycolysis^9, 10^. Additionally, microglia are also involved in maintaining brain lipid and lipoprotein homeostasis, a process that is often disrupted in disease, leading to the accumulation of lipid droplets (LDs)^11, 12^ ^13^. As a result of these metabolic adaptations, microglia undergo morphological changes. While the precise functional profiles of the different morphological states of microglia remain unclear, it is generally recognized that activated microglia alter their size and shape as they transition between homeostatic and disease-associated functions^7,14^. However, this phenotypic switching is largely context dependent. For example, microglia exhibit a hyper-ramified, highly complex phenotype in models of chronic stress^15^ and a rod-like and ameboid phenotype in models of neurodegenerative disease and aging^16–18^. Additionally, sex differences in microglia also dictate their morphology and function, adding another layer of complexity when assigning their shape to their role in the brain^16, 19^. Therefore, tools to study the cellular, molecular, and genetic factors that drive disease-associated metabolic and morphological changes in microglia are highly sought, not only to define disease mechanisms but to investigate novel interventions that restore microglial function and improve disease outcomes.

While *in vivo* models are useful for observing microglial changes in a physiological context, *in vitro* platforms are essential for investigating cellular mechanisms that are not easily testable *in vivo*. There are many *in vitro* systems to study changes in microglia activation and metabolic states (e.g., iPSC-derived microglia, immortalized microglia, and primary microglia (pMG)). However, each method presents its own strengths and limitations. For example, while iPSC-derived microglia are capable of reflecting age or disease-associated states^20^, they are expensive to maintain and exhibit genetic variability that can impact their reproducibility^21^. Comparatively, immortalized cell lines (i.e., HMC3 and BV-2) provide a less expensive and more homogenous platform for studying microglia. However, immortalized microglia do not accurately reflect *in vivo* microglia states and have been reported to be less responsive to external stimuli compared to pMG^22^. While both iPSC-derived and immortalized microglia have their individual strengths, pMG remain increasingly valuable when using transgenic mice to understand how specific genes regulate microglial function to protect against or promote disease. The ability to extract microglia from these complex models to empirically determine cellular and molecular mechanisms is crucial to expanding our understanding of disease processes. Despite the understanding that pMG may not reflect disease-related processes as accurately as iPSC-derived microglia, they allow us to utilize powerful genetic models to better reflect *in vivo* cell states than immortalized cells and offer a more financially accessible option^22, 23^.

Although primary microglia can be isolated using FACS or MAG sorting^24^, long sorting processes can potentially disrupt microglia activation status and, consequently, microglia metabolism^25^. Therefore, “shake-off” methods from mixed glial cultures are the quickest way to isolate a relatively pure, undisrupted microglia population for subsequent experiments. These methods have been used for decades and have been described previously^23, 26^. In brief, isolating microglia from a mixed-glial culture involves plating a whole-brain single-cell suspension, then shaking the flask to dissociate the loosely adherent microglia from the top layer of the mixed glial culture. However, this method is limited by the timing of available mice and the scalability of cell preparations. Although previous studies have indicated that microglia can be frozen down without impairing their function^27, 28^, no clear methods or functional assessment exist for pMG from frozen, whole-brain, mixed glial cultures. Similarly, multiple microglial “shakes” from a single mixed glial culture can introduce variability between experiments, but the impact of shake number on microglial function has never been formally reported. A deeper understanding of microglia shake-off protocols is critical for researchers studying microglia and neurological diseases, as these methods would allow precise temporal control over culture timing, facilitate sample banking, and enable scalable experimentation, all while maintaining the overall functional and metabolic integrity of the microglia.

Here, we established a protocol for freezing mixed glial cultures to enable banking for future microglia isolations. We also examined whether pMG isolated from the first or second shake exhibited comparable morphological and metabolic changes in response to various environmental stimuli across fresh (FS) and frozen (FZ) conditions. Although freezing introduced measurable differences in microglial morphology and LD accumulation, microglia derived from frozen cultures responded similarly to environmental challenges, such as high-glucose and LPS, as those from fresh cultures. Therefore, while FZ primary microglia recapitulate functional responses, maintaining consistency in the use of either FS or FZ cells within an experiment is strongly recommended.

## Results

### Frozen murine mixed glial cultures can be thawed and shaken to isolate pMG

Previous literature suggests that primary mixed glial cultures can be frozen down for future use^27, 28^, but does not describe freeze-thaw protocols in detail or compare microglia functionality after freezing. Nonetheless, these resources served as the starting point for optimizing and validating the freeze-thaw protocol for mixed glial cultures described in the present study.

In brief, the brains of post-natal day 0-2 (p0-p2) mouse pups were isolated and kept in Hanks’ Balanced Salt Solution (HBSS) on ice until the meninges were successfully removed. The brains were then pooled and incubated with a digestion buffer to generate a single-cell solution. Each brain was split between two 25 cm^2^ flasks (i.e., half a brain per flask) to generate a mixed glial culture (Figure 1A). The cultures were grown in primary media containing Dulbecco’s Modified Eagle Medium/Nutrient Mixture F-12 (DMEM/F12), 10% fetal bovine serum (FBS), and 1% penicillin/streptomycin (P/S). Although the mixed glial cultures had several pieces of debris attached to the plate after isolation due to the nature of this preparation, the debris gradually detached after the first full media change (Figure 1B).

**Figure 1:**
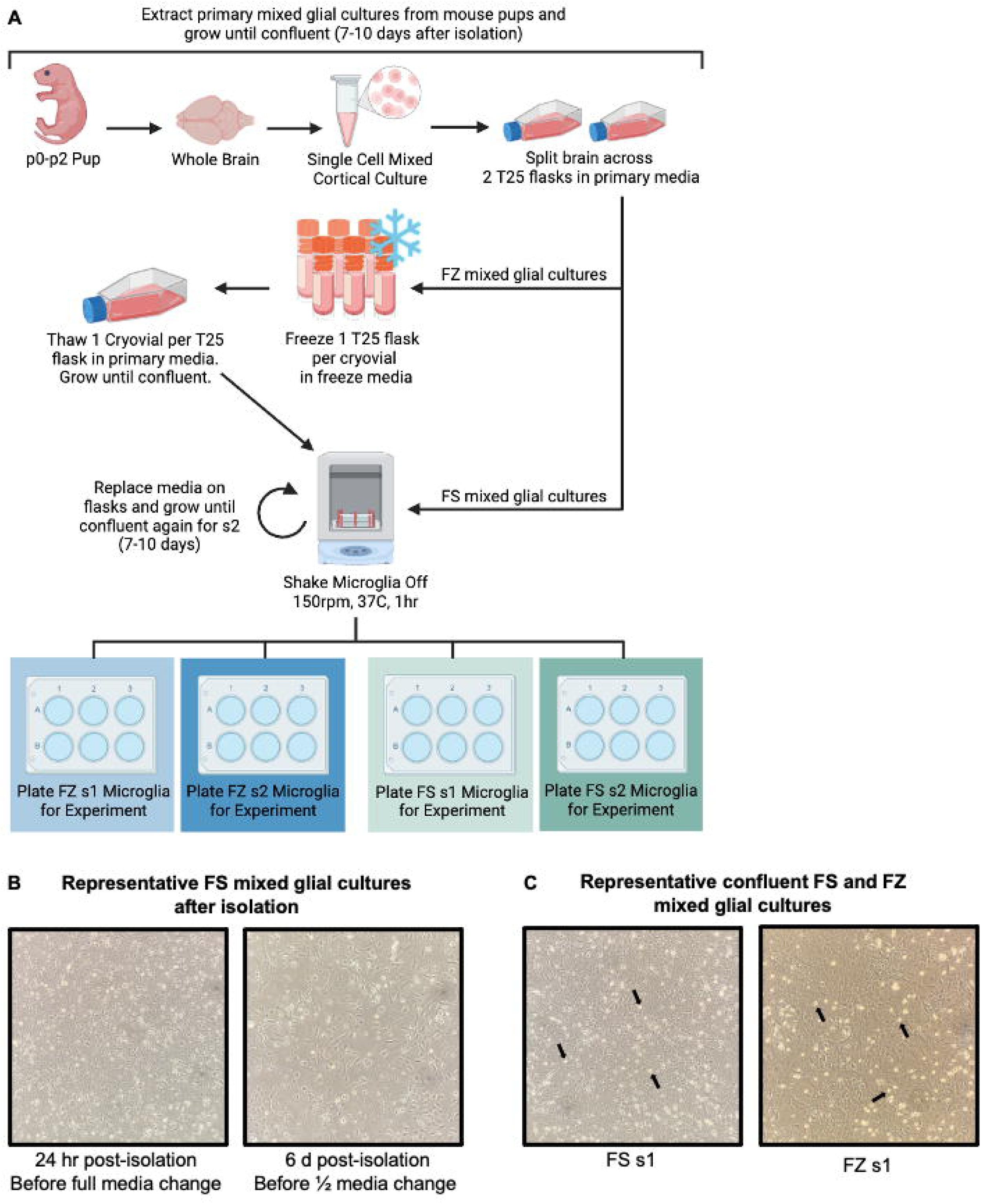
pMG can be isolated from both fresh and previously frozen mixed glial cultures. **(A)** Workflow for the extraction and freeze-thaw of mixed glial cultures from p0-p2 mouse pups as well as the isolation of primary microglia (pMG) from fresh (FS) and frozen (FZ) mixed glial cultures. **(B)** Representative images of FS mixed glial cultures at 10X after extraction from p0-p2 pups, but before reaching maximum confluency for freeze down or pMG shake. **(C)** Representative images of confluent FS and FZ mixed glial cultures at 10X immediately before shake 1 (s1) illustrating the similarities between the two. Black arrows indicate examples of small, circular, bright microglia floating in the media.

To generate FS shake 1 (s1) microglia cultures, microglia were isolated once maximum confluency was reached after about 7-10 days (Figure 1C). At this stage, the mixed glial culture began to produce small, circular, bright microglia floating in the media and loosely attached to the plate (Figure 1C). Maximum confluency was defined as when microglia accounted for approximately 20% of the cells in each field of view (Figure 1C). Microglia were isolated from the mixed glial culture using a shake-off method, where flasks were put on a shaker for 1 hour at 150 rpm and 37°C. Because this shaker lacked CO_2_ regulation, the flask filters were wrapped tightly with parafilm to prevent CO_2_ loss. This method has been reliably used by our lab and others to release solely microglia from the surface of the plate for isolation^23, 29, 30^. Microglia can then be isolated by centrifuging the media at 500 xg for 5 minutes and pelleting the cells. pMG were then resuspended and seeded at a density of 130,000 cells per well in a 24-well plate (Figure 1A). Proper density of pMG plating is critical to ensure that cells are in close proximity to maintain the necessary cell-to-cell communication for optimal growth and health. This is especially important for this protocol, given that we do not supplement the microglia culture media with additional growth factors or cytokines. After s1, mixed glial cultures received fresh media and were left to grow to maximum confluency again for approximately 7-10 days, producing FS shake 2 (s2) microglia (Figure 1A). These cells were isolated and plated identically to s1 microglia. We did not notice any difference in the average microglia yield from FS s1 (∼165k per flask) and FS s2 cultures (∼148k per flask), consistent with previous literature^26^.

To generate FZ cultures, mixed glial cultures were frozen down once they had fully tiled the surface of the flask, but before they reached maximum confluency with floating microglia, approximately 7 days after isolation (Figure 1A). Mixed glial cultures were removed from the surface of the flask using trypsin, pelleted, and resuspended in fresh primary media. Cells were resuspended with 100 μL of media for every flask frozen down, with one flask ultimately banked per cryovial. For example, if each cryovial were to receive 100 μL of cell solution and 10 flasks were being frozen down, then the cells would be resuspended in 1mL of media. In early trials of this protocol, we attempted to split each confluent flask between two cryovials (e.g., ½ flask per cryovial). However, decreasing the amount of cells per vial coupled with natural cell loss during the freeze-thaw process resulted in a much slower recovery and perceived proliferation after plating (visualized as a reduced number of cells) than when each 25 cm^2^ flask was cryopreserved in a single cryovial (Figure S1). Therefore, we recommend freezing each confluent 25 cm^2^ flask of mixed glial culture as a single cryovial.

After resuspension, 100 μL of cell solution was added to each cryovial, quickly followed by 900 μL of freeze media. This order is crucial to prevent the cells from being exposed to excessive amounts of dimethyl sulfoxide (DMSO) for prolonged periods, as freeze media for these experiments consisted of 70% primary media, 20% FBS, and 10% (DMSO). However, because the freeze media was diluted with the cell suspension in primary media, the final concentrations in the cryovial were 73% primary media, 18% FBS, and 9% DMSO. We also tried a freeze media combination of 90% FBS and 10% DMSO to determine if a greater FBS concentration would increase cell viability, although we noted no distinct differences (Figure S1). Cryovials were quickly placed in a freezing container with isopropanol and stored overnight in a -80°C freezer, then moved to liquid nitrogen (LN2) the next morning.

After being frozen for 4 days, FZ mixed glial cultures were thawed and plated on 25 cm^2^ flasks, assuming one flask per cryovial. While we did not directly test how long mixed glial cultures were viable after storage at -80°C, we had kept them frozen for 4 weeks in early trials without issue and anticipate that they can be cryopreserved in LN2 long-term. Thawed cells were grown until maximum confluency was reached, typically about 10-14 days. FZ s1 microglia were then isolated using the shake-off procedure and plated as previously described. Mixed glial cultures received fresh media and were left to grow to maximum confluency again, producing FZ s2 microglia. These cells were isolated and plated identically to s1 microglia. While we did not identify any differences in the average number of microglia isolated between all FS cultures and FZ s1 cultures (∼162k per flask), we did notice lower yields produced by FZ s2 cultures (∼100k per flask).

Once cells were adhered to the plate, they were treated for 24 hours with either control media (17.5 mM glucose), high-glucose media (High Glu, 35 mM), or lipopolysaccharide (LPS) media (LPS, 10 ng/mL LPS + 17.5 mM glucose) for 24 hours. The goal with these environmental stimuli was to prompt the microglia to respond by changing either their activation status or metabolism. Therefore, the concentrations were chosen to be sufficient to induce change rather than fully disrupting the cells or risking cytotoxicity, as described previously^31^.

After treatment, pMG were fixed with 4% paraformaldehyde (PFA) and stained with phalloidin (Ph), an F-actin stain that targets the entire cell^32^, and BODIPY, a neutral lipid stain that targets lipid droplets^33^. CellProfiler image analysis software was used to quantify changes in cell morphology^34^, using Ph, and LD accumulation, using BODIPY, in response to external stimuli (Figure S2).

### pMG isolated from FS and FZ mixed glial cultures undergo similar morphological responses to environmental stimuli

Ph-positive cells were used to quantify changes in cell morphology, which can be used as a proxy for microglial activation status. Although pMG *in vitro* do not perfectly recapitulate the same responses to stimuli seen *in vivo*, changes in microglial responsiveness *in vitro* can be garnered from changes in cell size and shape^14, 35^. Changes in microglia size were assessed using total Ph area and maximum radius (max radius) measurements. Generally, increases in cell size are associated with microglia activation, as seen in response to LPS treatment^36^ and pathological stimuli^37^. While morphology does not directly indicate function, enlarged cell bodies are typically associated with increased phagocytosis and migration towards inflammation^7^.

Ph staining allowed the visualization of cell morphology following High Glu and LPS treatment. Overall, pMG exhibited few, but limited, changes in response to High Glu conditions. Although pMG remained small and somewhat ameboid in both conditions, these changes are consistent across FS and FZ cultures of both shakes (Figure 2). In response to LPS, pMG changed shape and size, appearing larger and less circular than in control and High Glu-treated cells (Figure 2). Again, these qualitative morphological changes are consistent between FS and FZ cultures of both shakes (Figure 2).

**Figure 2:**
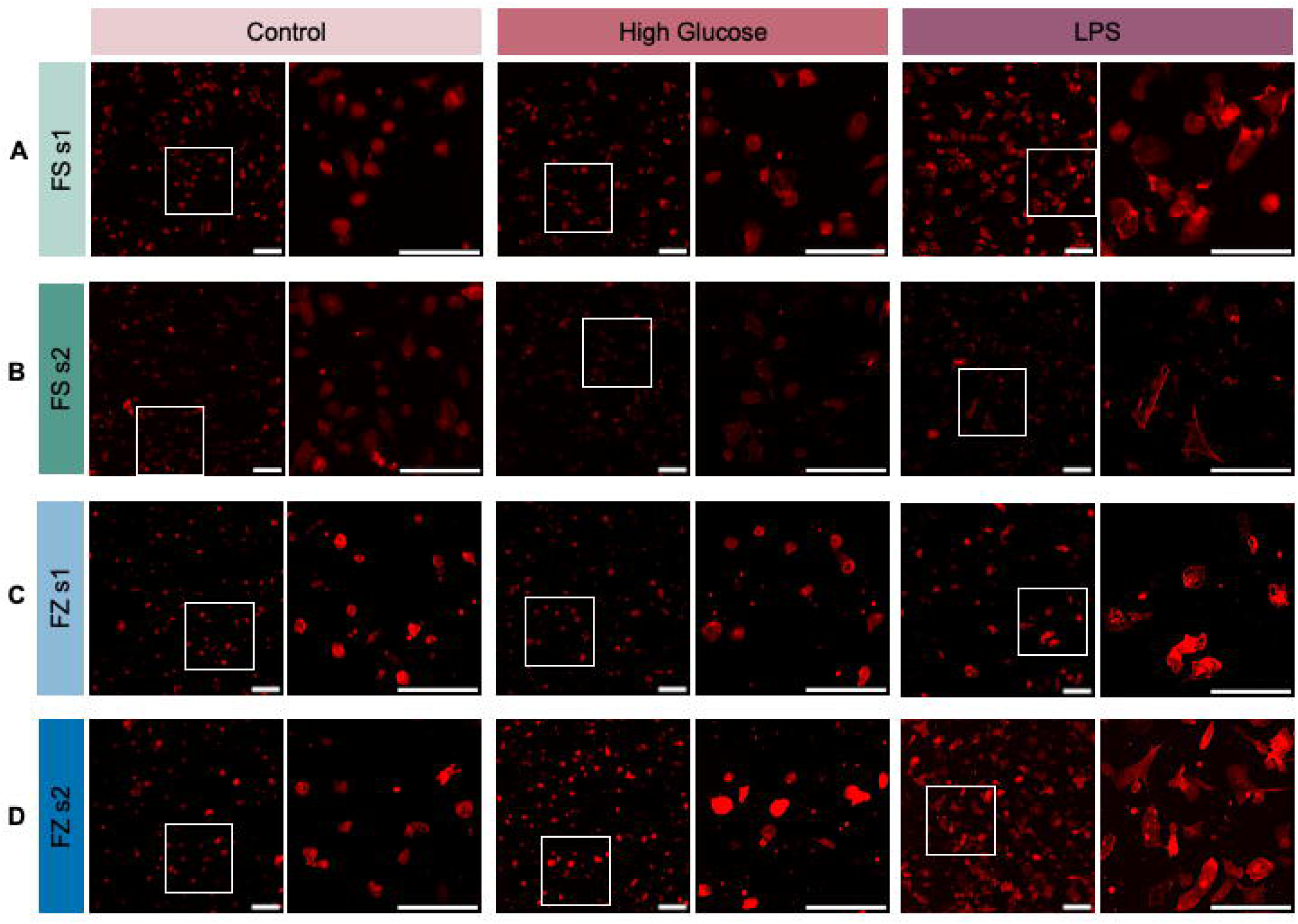
Representative images of phalloidin (Ph)-stained primary microglia (pMG). Changes in pMG shape and size in response to control, high glucose (High Glu), and lipopolysaccharide (LPS) from **(A)** fresh (FS) shake 1 (s1), **(B)** FS shake 2 (s2), **(C)** frozen (FZ) s1, and **(D)** FZ s2 cultures. White boxes indicate the region selected for the zoomed-in image seen to the right. Scale bars represent 100 µm.

Our quantifications of Ph staining size corroborated our qualitative observations. Specifically, we found that FS and FZ pMG from both shakes showed no changes in Ph area when grown in High Glu conditions (Figure 3a). While FS s1, FS s2, and FZ s1 High Glu microglia also showed no differences in maximum radius, FZ s2 microglia displayed a slight decrease, suggesting that changes in cell size in response to High Glu in different shakes may need to be investigated more carefully (Figure 3b). In contrast, following LPS treatment FS and FZ microglia from both shakes displayed a consistent increase in Ph area and max radius (Figure 3a-b).

**Figure 3:**
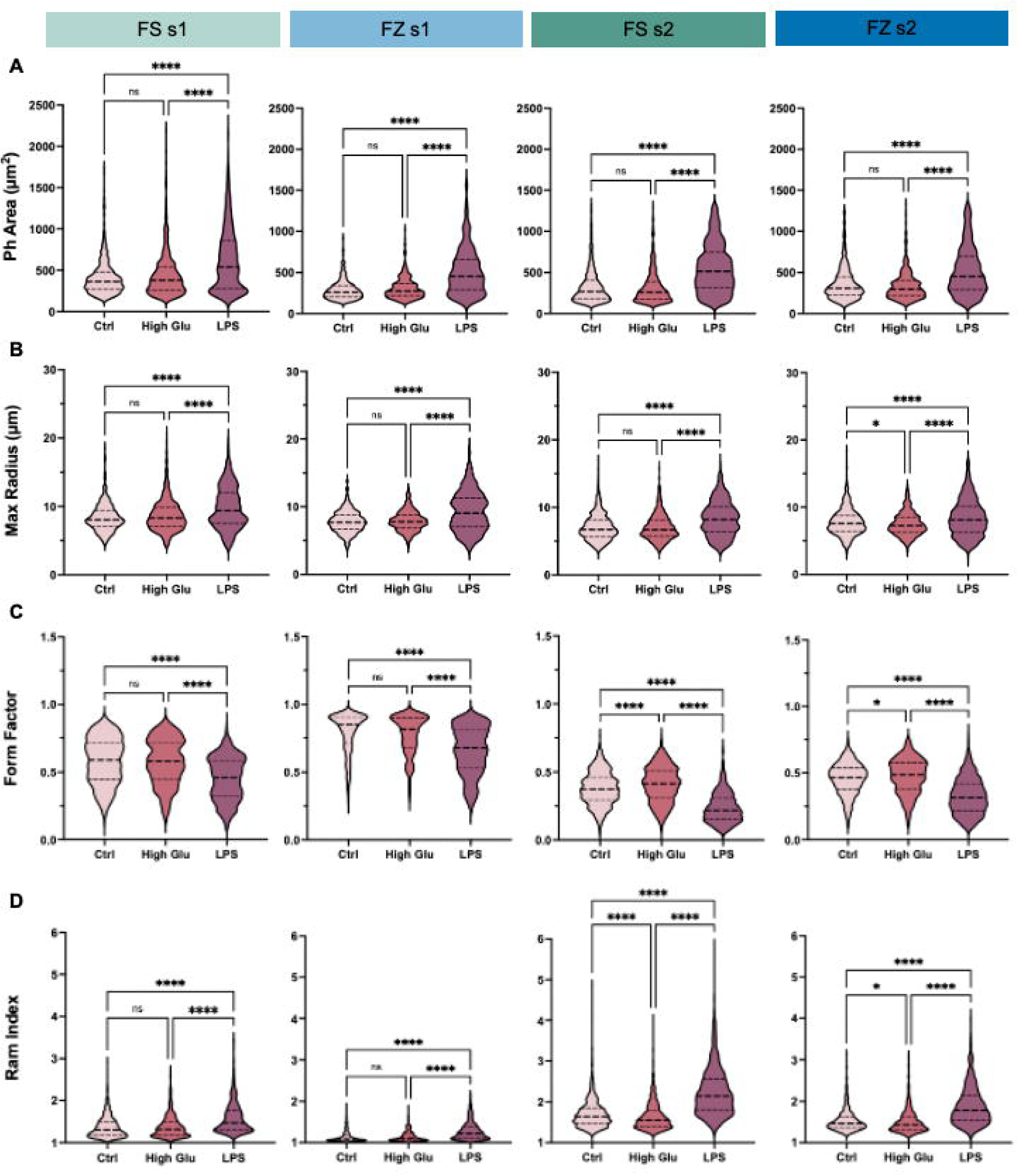
Changes in shape and size do not differ between primary microglia (pMG) isolated from fresh (FS) and frozen (FZ) mixed glial cultures in response to high glucose (High Glu) and lipopolysaccharide (LPS) stimulation. Changes in pMG size were quantified using **(A) Phalloidin (**Ph) area (µm^2^) and **(B)** max radius (µm). Changes in pMG shape were quantified using **(C)** form factor and **(D)** ramification (ram) index. Data are presented as quartiles (n = 300-1,500 cells per group), *P < 0.05, ****P < 0.0001 (Kruskal-Wallis nonparametric test with Dunn’s multiple comparisons test), ns = not significant. See also Figure S3.

Differences in cell shape were measured by changes in the complexity of the cell, which considers both area and perimeter measurements to assess if the cell is more circular or highly processed. Complexity was measured using form factor ((4*π*area)/perimeter^2^) and ramification index (ram index, (perimeter/area)/(2(π/area)^1/2^)). High complexity is indicated by a low form factor (form factor of a circle = 1) and a high ramification index. While microglia shape will vary in response to different stimuli, cells will generally increase in complexity in response to stress^38, 39^ or pro-inflammatory cytokines^40^, which are associated with increased synaptic modifications^38, 41^. pMG showed no change in form factor or ram index in response to High Glu in FS and FZ s1 cultures (Figure 3c-d). In comparison, in FS and FZ s2 cultures, High Glu microglia displayed a slight increase in form factor and a decrease in ramification index, suggesting that the impacts of High Glu on microglia shape and complexity are shake-specific and not influenced by FS-FZ status (Figure 3c-d). Comparatively, microglia treated with LPS displayed significant decreases in form factor and increases in ramification index in FS and FZ conditions in both shakes, indicating consistently increased complexity regardless of conditions (Figure 3c-d).

Despite parallels in the magnitude and direction of differences from controls in High Glu and LPS-treated microglia across FS-FZ and shake status, comparisons within each group revealed differences in the absolute measurements across stimuli. Specifically, FS s1, FS s2, FZ s1, and FZ s2 had significantly different morphological metrics in control, High Glu, and LPS treatments (Figure S3).

Despite this, measurements for each group are within a relatively similar range, suggesting that the differences between FS, FZ, s1, and s2 are minimal (Figure S3). However, since morphology does change across treatments, this data suggests that absolute measurements from pMG at different FS-FZ statuses or shakes should not be used interchangeably or compared within a given experiment. Given the robust response to LPS, we would recommend considering the change from baseline as a useful metric for data spanning several experiments if cell availability is limited. Still, the overall consistent changes in cell shape and size between pMG from FS and FZ cultures suggest that FZ-derived pMG can be reliably used to assess microglia size and shape as a proxy for activation status in response to environmental stimuli.

### Environmental stimuli similarly modulate LD accumulation in pMG from FS and FZ mixed glial cultures

pMG BODIPY staining was used to quantify changes in LD accumulation, which can be used as a proxy for microglial immune dysregulation and activation^13, 42^. LD accumulation is associated with microglial dysfunction, as it leads to decreased phagocytosis and increased pro-inflammatory cytokine release^13, 42, 43^. LD accumulation is thought to be a consequence of inflammatory polarization, which is linked to a metabolic shift towards glycolysis, and the increased production of intermediary metabolites that drive *de novo* lipogenesis and LD accumulation^42^. Increased fatty acid synthesis, coupled with decreased fatty acid breakdown, is associated with increased secretion of cytokines and decreased ATP production, which dampens the cell’s phagocytic abilities^13, 42^.

BODIPY-positive cells were observed in all conditions (Figure 4a-d). Overall, pMG did not show large changes in BODIPY staining in High Glu conditions, which was consistent across FS-FZ status and shake status. However, in response to LPS treatment, pMG appeared to have smaller amounts of BODIPY-positive area per Ph-positive cell (Figure 4a-d). Notably, this trend was consistent across FS-FZ status in both shakes.

**Figure 4:**
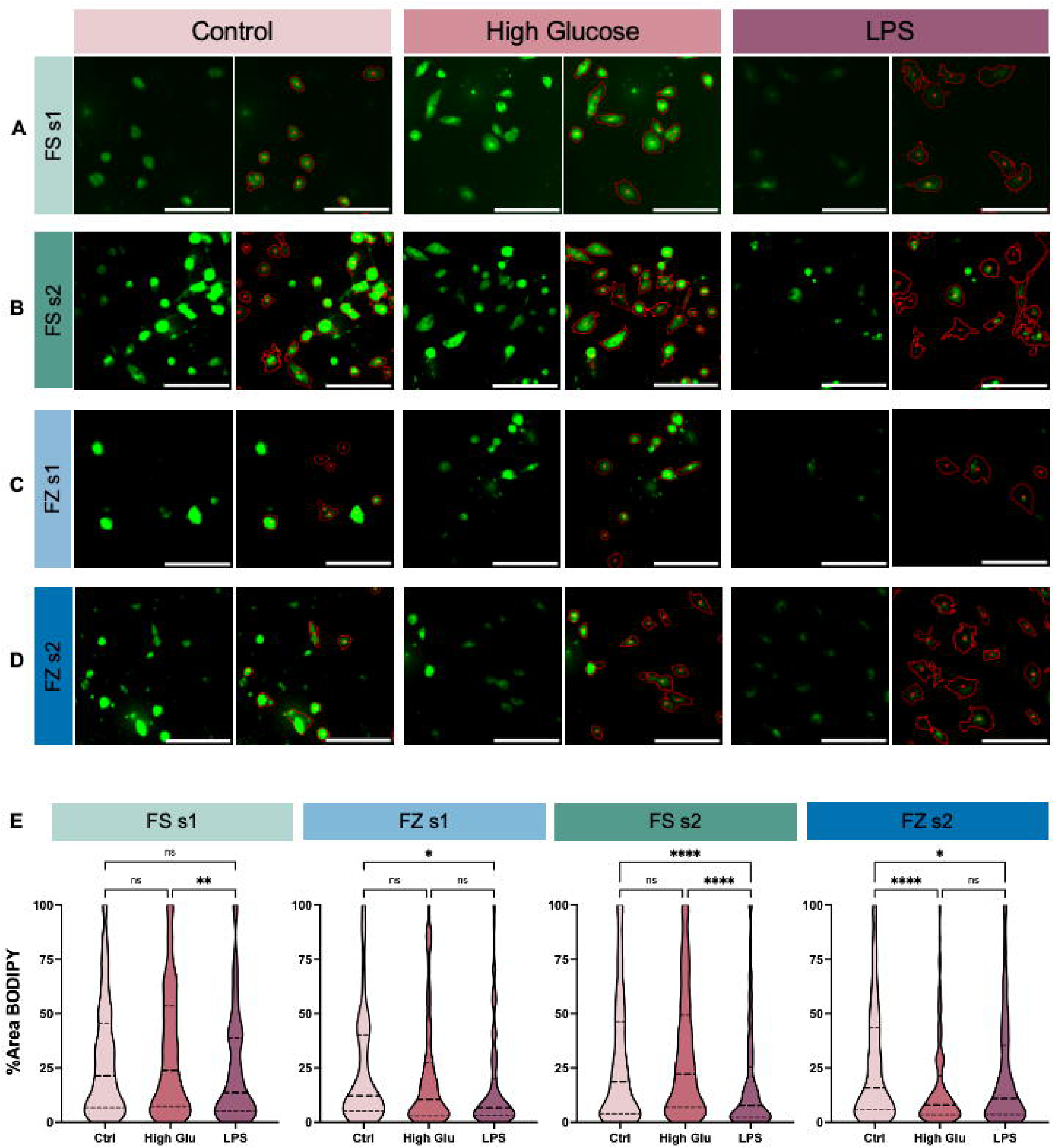
Changes in BODIPY-stained lipid droplet (LD) accumulation do not differ between pMG isolated from fresh (FS) and frozen (FZ) mixed glial cultures in response to high glucose (High Glu) and lipopolysaccharide (LPS) stimulation. Representative images of BODIPY-positive regions within Ph cell masks illustrating changes in primary microglia (pMG) in response to control, High Glu, and LPS from **(A)** FS shake (s1), **(B)** FS shake 2 (s2), **(C)** FZ s1, and **(D)** FZ s2 cultures. **(E)** Changes in pMG LD accumulation were quantified by calculating the percent area of the cell covered by BODIPY. Scale bars represent 100 µm. Data are presented as quartiles (n = 8-9 images per group), *P < 0.05, **P < 0.01 ****P < 0.0001 (Kruskal-Wallis nonparametric test with Dunn’s multiple comparisons test), ns = not significant. See also Figure S4.

To investigate our qualitative observations, changes in LD accumulation were measured by the percentage of each cell area covered by BODIPY as well as the percentage of BODIPY-positive cells per image. Microglia from FS s1, FS s2, and FZ s1 treated with High Glu displayed no differences in the percent area covered by BODIPY, while FZ s2 microglia displayed a significant decrease (Figure 4e). In response to LPS, FS s2, FZ s1, and FZ s2 showed decreases in the percent area covered by BODIPY, whereas FS s1 cultures showed no significant change, despite a clear decreasing trend that mimics the changes observed in the other conditions (Figure 4e). Again, this suggests that while pMG from FS and FZ cultures largely respond similarly to stimuli, there may be shake-specific differences. Furthermore, microglia from each condition showed no significant differences in the percentage of BODIPY-positive cells in response to High Glu or LPS (Figure S4a).

Similar to our morphological analysis, we also investigated differences in the absolute measurements of BODIPY across each stimulus. Importantly, there are no differences in percent area covered by BODIPY in control conditions across FS-FZ status or shake number, indicating that all cells are starting at similar metabolic states regardless of whether they were frozen or shaken multiple times (Figure S4b). However, there is a condition-specific effect with High Glu and LPS treatment (Figure S4b). This suggests that while the direction of change in percent area covered by BODIPY is consistent across conditions, the total percent area covered by BODIPY after treatment may vary. Despite this, the percentage of BODIPY-positive cells is not significantly different across FS-FZ status or shakes after control, High Glu, or LPS treatments, with notably high variability between conditions (Figure S4c). In sum, while there may be shake-specific effects, these results suggest that pMG from FS and FZ mixed glial cultures have consistent changes in LD accumulation, and therefore metabolism, in response to environmental stimuli.

## Discussion

The ability to freeze down mixed glial cultures offers previously unattainable control over pMG cells. In this paper, we show that it is possible to produce responsive microglia from previously frozen primary mixed glial cultures. We tested this by treating s1 and s2 pMG from both FS and FZ cultures with High Glu or LPS. To assess their responsiveness to external stimuli, we measured microglia morphology and LD accumulation as a proxy for their activation status and associated metabolic changes. In response to LPS, we identified that pMG consistently increased in size and complexity across FS-FZ status and shake number (Figure 2-3). Additionally, LPS induced consistent reductions in the percent area of the cell covered by BODIPY in both FS and FZ cultures (Figure 4). In response to High Glu, we identified no changes in pMG circularity, complexity, or LD accumulation (Figure 2-4). This observation is also consistent across shake and FS-FZ status (Figure 2-4). Overall, our results indicate that pMG from FS and FZ have similar responses in size, complexity, and percent area covered by BODIPY in response to external stimuli. However, there may be variations between shake numbers. Hence, for the first time, we report a validated method for freezing primary mixed glial cultures to yield pMG that respond to external stimuli in the same way as cells isolated from fresh mixed glial cultures.

We show that pMG isolated from both FS and FZ cultures become larger in response to LPS exposure, a phenomenon which is consistent with previous reports of LPS broadly promoting microglial activation^14, 36^. In contrast, LPS also reduced LD area and increased cell complexity, which is perhaps unexpected given the relationship between LPS and microglial activation^13, 42, 44^. However, since microglial activation exists on a spectrum, it is possible that the LPS dose (10 ng/mL) used in this study was sufficient to induce microglial activation without causing metabolic reprogramming and microglial dysfunction. This is supported by the shift from amoeboid to elongated morphology observed in pMG from both FS and FZ cultures treated with LPS (Figure 3). In further support, previous studies linking LPS to microglial metabolic reprogramming and LD accumulation have used much higher doses of LPS 1-5 μg/mL^13, 45, 46^. Therefore, we postulate that our LPS model induces a response akin to microglial priming, which occurs when they receive a low dose of external stimuli that trains them to produce a heightened response to subsequent immune challenges^47–49^. While priming is on the same spectrum of microglia activation, it is morphologically and functionally distinct^48^. Therefore, we propose that because these pMG have not been cultured with cytokines and are naïve to inflammatory stimuli, a single dose of LPS may be sufficient to induce a priming-like effect that exists on the spectrum of activation. In comparable *in vitro* models, primed pMG exhibit a ramified, complex morphology, which parallels our results and is distinct from the ameboid shape expected in activated microglia^48^. While the exact impacts of microglia exposure to low concentrations of LPS on LD accumulation are unknown, primed pMG are also associated with decreased cytokine release^48^. Given the known association between LD accumulation and increased cytokine release^13, 42^, we postulate that the observed decrease in LD accumulation following LPS is indicative of a priming–like response in these naïve pMG. Additionally, due to our inability to resolve individual LD dynamics from our images, it is possible that this initial response to low-dose LPS induces changes that are more nuanced than BODIPY area and number of BODIPY-positive cells.

Previous literature has described that high glucose conditions are sufficient to induce an ameboid morphology with minimal complexity^50, 51^. Our inability to replicate this pattern in our High Glu treatment may be due to the pMG culture media, which already contains a high glucose concentration (e.g., 17.5mM). Since diabetes patients with elevated cerebrospinal fluid (CSF) glucose measurements only reach ∼10 mM glucose^52^, the control levels of glucose may be too high to resolve any differences from High Glu-treated microglia. In fact, recent work that describes these morphological shifts in response to High Glu treatment either uses significantly higher, non-physiologically relevant concentrations in the High Glu condition^51, 53^ or significantly lower amounts in the control conditions^50^. Unforeseen impacts of control levels of glucose may also explain the lack of expected LD accumulation seen between control and High Glu-treated pMG^54^. However, similar to our observations in LPS-treated pMG, we may be inducing changes in LD dynamics at glucose concentrations between 17.5 mM and 35 mM that cannot be resolved with our current imaging platform.

In summary, we see consistent microglial morphological and metabolic changes in pMG in response to stimuli between FS and FZ cultures. Despite these similarities, differences in responsiveness across conditions suggest that absolute metrics from microglia in different culture conditions (FS-FZ status and shake number) should not be directly compared. Rather, we recommend consistently using either FS or FZ cells of a specific shake throughout a given experiment. Overall, we report that mixed glial cultures can be successfully frozen to yield typically responsive microglia for downstream analysis.

### Study Limitations

While producing responsive microglia from frozen cultures is a major advance in the field of *in vitro* methodology, this paper is not without its limitations. Our outputs of cell morphology and LD accumulation as metrics of microglia activation status and metabolism broadly assess the responsiveness of microglia to external stimuli. However, several additional metrics could be added to quantify more nuanced changes in microglia function (i.e., phagocytosis, cytokine release, and Seahorse metabolic assays). Similarly, transcriptomics analysis could be performed on microglia from FS and FZ cultures from both shakes to investigate more robust changes in their inflammatory profile as a result of the freeze-thaw process. However, this is beyond the scope of this study, as we aimed to only establish a method to freeze and revive viable mixed glial cultures for future use. Additionally, we did not determine how long mixed glial cultures can remain frozen before thawing to generate responsive microglia. Future studies can ensure that long-term storage of mixed glial cultures in LN2 is viable. Similarly, future studies can also investigate the use of this protocol for transgenic mice to determine whether the inclusion of a Cre-recombinase transgene or constitutive knock-out impacts the viability of mixed glial cultures when frozen. Lastly, we only tested the response of microglia to 35 mM High Glu and 10 ng/mL LPS conditions. Therefore, we are not able to conclude whether there is a stimulus- or concentration-specific effect that may differ between fresh and frozen cultures. As a result, we recommend that researchers using primary microglia from frozen mixed glial cultures investigate changes in their desired output following treatment of interest between fresh and frozen cultures to ensure there are no shake- or treatment-specific effects.

## Acknowledgements

We thank Dr. Wendy Macklin and the members of the Macklin lab for sharing their incubated shaker with us. We also thank the University of Colorado Anschutz Medical Campus Division of Endocrinology, Diabetes, and Metabolism for their continued financial support of the BioSpa Cytation 5 Imager. This work was supported by NIH-NIA R21AG091650 awarded to KDB, T3263032819 awarded to AMS, and T32DK120521 awarded to JEH.

## Author Contributions

AMS and KDB conceived the project. AMS designed, performed, and analyzed experiments. AMS and KDB wrote and revised the manuscript with contributions from all authors. KMB and GJP assisted with performing experiments. JEH assisted with developing the CellProfiler analysis pipeline.

## Declaration of Interests

The authors have no competing interests to declare.

## Resource Availability

Lead contact: Requests for further information and resources should be directed to and will be fulfilled by the lead contact, Kimberley Bruce, PhD.

Materials availability: This study did not generate any new unique reagents.

Data and code availability: All imaging pipelines have been deposited on GitHub and are publicly available at https://github.com/asturno/CellProfiler as of the date of publication.

## Declaration of generative AI and AI-assisted technologies in the writing process

During the preparation of this work, the authors used Perplexity and Copilot to proofread portions of the document for clarity. After using this tool, the authors reviewed and edited the content as needed and take full responsibility for the content of the publication.

## Methods

### Mixed Glial Culture Isolation from p0-p2 Mouse Pups

All investigations in this study were performed using C57BL/6J mice on protocols (Ref #0115 and #0815) that were approved by the University of Colorado Institutional Animal Care and Use Committee (IACUC), ensuring compliance with the recommendations in the Guide for the Care and Use of Laboratory Animals, Animal Welfare Act and PHS Policy. Mice were housed no more than 5 mice per cage and maintained at ∼ 20 °C with a 14-10 hour light/dark cycle and given unrestricted access to a standard laboratory diet (Diet 8640; Harlan Teklad, Madison, WI, USA) and water. Mice were bred in duos, and pups that were not isolated were weaned at 28 days.

Male and female mouse pups from p0-p1 were pooled together and anesthetized on ice until they no longer responded to a toe pinch. Pups were decapitated quickly with scissors, then the brains were quickly removed and placed in a petri dish with cold Hanks’ Balanced Salt Solution (HBSS, Gibco) on ice. Brains were dissected in groups of five to avoid them sitting on ice for too long. If more than five pups needed to be isolated in a day, we would wait until the first group was done to begin anesthetizing and removing the brains of the next group.

After dissection, the brain was placed in cold HBSS in a fresh petri dish under the dissection scope to remove all meninges using forceps. The whole brain without meninges was placed into a 6-well plate (Corning) containing HBSS on ice. Up to 3 brains were pooled into each well of a 6-well plate.

After all dissections were complete, the 6-well plate containing the brains was moved into the cell culture hood. The HBSS was carefully aspirated, and 1 mL of 37 °C trypsin (0.05%, Gibco) per brain was added to each well for dissociation. Brains were minced with fine forceps and placed in the incubator (37 °C, 5% CO_2_). After 10 minutes, the plate was removed and gently swirled to mix the brains in the trypsin and increase dissociation, then placed back in the incubator for another 10 minutes. After the full 20 minutes, 3 mL of warmed primary media (DMEM/F12 50/50 1X (Gibco) + 10% FBS (GeminiBio) + 1% P/S (Corning)) was added to each well to stop the trypsin reaction. Each well was triturated 10x with a P1000 pipette and transferred into a 15 mL conical tube. The tubes were left undisturbed for 2-3 minutes to allow large debris to settle. Avoiding this large debris, the supernatant was filtered through a 70 μm filter, pre-wet with primary media, into a 50 mL conical tube. 3 mL of fresh media was added back to the debris to triturate 10X with a P1000 pipette again before filtering into the same 50 mL conical. The filtered single cell suspension was centrifuged at 500 xg for 5 minutes, and the supernatant was aspirated. The pellet was resuspended in the appropriate volume of media to seed each brain into two T25 flasks at 4 mL/flask. 4mL of resuspended cell solution was gently pipetted onto a PDL-coated (Gibco, 0.05 mg/mL) 25 cm^2^ flask (Corning). PDL was incubated on the flask for a minimum of 1 hour, then rinsed twice with PBS (Cytiva) before cells were added.

After being incubated for 24 hours at 37 °C with 5% CO_2_, the cells were fully adhered. Existing media was aspirated, and flasks were rinsed twice with warmed primary media to remove debris before adding back 4 mL of fresh media. Following this, half media changes were performed every 3-4 days (typically every Monday and Thursday or Tuesday and Friday). Flasks were tilted to pool all media in the bottle corner and allow floating cells to sink to the bottom. 2 mL of media was removed from the top of the pool to avoid any floating cells. A fresh 2 mL was then added back into the flask to bring the final volume back up to 4 mL.

### Primary Microglia Isolation from Mixed Glial Culture

Flasks with mixed glial cultures were grown until they were confluent and had significant granularity (small, bright dots that are likely floating microglia). To isolate microglia from the mixed glial culture, the filters and caps of the flasks were tightly parafilmed before placing them in an incubated shaker at 150 rpm for 1 hour. Following the shake, the media containing detached microglia was removed with a serological pipette and added to 15 mL conical tubes. Each flask was rinsed thoroughly with 2 mL of media, which was then added to the same conical. If the flask was being saved for a second shake, a fresh 4 mL of media was added, and the flasks were put back in the incubator. Meanwhile, the conical tubes containing microglia were centrifuged at 500 xg for 5 minutes, then resuspended in 1mL of media.

### Plating Primary Microglia

Once microglia were isolated from the mixed glial culture, cells were counted using a cell counter (Countess) to prepare for plating. This was achieved by mixing 10 μL of cell solution and 10 μL of trypan blue (Gibco), then adding 10 μL of this mixture to the cell counter cassette. Using the cell count, cells were resuspended in primary media to achieve roughly 130,000 cells in 500 μL per well. For all immunohistochemistry and imaging experiments, 500 μL of the resuspended cell solution was added to each 1.54 cm^2^ well of a 24-well plate (Ibidi). Before any treatment, cells were incubated for 4 hours to allow for adherence.

### Freezing Mixed Glial Cultures

To generate frozen microglia cultures, mixed glial cultures in 25 cm^2^ flasks were grown until confluent and granular, as described above. The media was aspirated from the cells, and each flask was rinsed twice with room temperature PBS. 1 mL of 37°C trypsin (Gibco, 0.05%) was added to fully cover each flask. After incubating cells in trypsin for 10 minutes, 3 mL of primary media was added to each flask to stop the reaction. The flasks were gently tapped to dislodge as many cells as possible, but still required scraping to fully detach all cells. Each flask was tilted, and all cells were washed down to one corner to be taken up and moved to a 15 mL conical tube and centrifuged at 500 xg for 5 minutes. The supernatant was aspirated, and cells were resuspended with 100 μL of media for every flask frozen down, with one flask ultimately banked per cryovial. For example, if each cryovial receives 100 μL of cell solution and 10 flasks were being frozen down, then the cell pellet would be resuspended in 1mL of media. 900 μL of freeze media (primary media + 20% FBS + 10% DMSO) was added to the cryotubes to bring each one to a final volume of 1 mL. However, because the freeze media was diluted with the cell suspension in primary media, the final concentrations were 73% primary media, 18% FBS, and 9% DMSO. All cryotubes were immediately placed in a room-temperature freezing container (Mr. Frosty, Thermo Scientific) containing isopropyl alcohol (Fisher), which was placed in a -80°C freezer. The following morning, all cryotubes were placed in LN2 for long-term storage.

### Thawing Frozen Mixed Glial Cultures

Cryotubes were thawed in a 37°C water bath until only a small ice chunk remained. Once thawed, an additional 1 mL of primary media was immediately added to each tube to dilute the DMSO. These 2 mLs were then transferred to a 15 mL conical tube. Each cryotube was rinsed with an additional 2 mL of media, which was then added to the same conical, which was spun at 500 xg for 5 minutes. During this time, PDL-coated flasks were rinsed twice with PBS. 4 mL of primary media was added to each 25 cm^2^ flask, and they were incubated to warm to about 37°C for cell plating. The cell pellet was resuspended in 1 mL of primary media and split across as many 25 cm^2^ flasks as cryotubes that were initially thawed (i.e., if 5 cryotubes were thawed, split the resuspended cell solution across 5 flasks).

Half media changes were performed every 3-4 days until full confluency and granularity were achieved, as described above.

### LPS and High Glucose Treatment of Primary Microglia Cells

Once confluent, n=3 1.54 cm^2^ wells of a 24-well plate were treated with 250 μL 3X concentrated lipopolysaccharide (LPS, Millipore Sigma) at 30 ng/mL. In culture, this makes the working 1X concentration 10 ng/mL and yields a final well volume of 750 μL . Similarly, n=3 1.54 cm^2^ of a 24 well-plate were treated with 250 μL of 3X concentrated sterile filtered D(+)-Glucose (ThermoScientific) at 105 mM. In culture, this makes the 1X working concentration 35 mM and yields a final well volume of 750 μL. Untreated primary media was used to dilute all LPS and glucose treatments and was also added to control wells to achieve a final volume of 750 μL. Cell treatments were added to existing media to avoid unnecessary disturbances to the cells, as they very easily detach from the plate. Treatments were left on cells for 24 hours in a 37°C incubator with 5% CO_2_.

After 24 hours of incubation with the treatment conditions, cells were slowly and gently rinsed once with PBS (Cytiva), then fixed with 4% PFA (Electron Microscopy Sciences) for 10 minutes at room temperature (RT). The cells were rinsed with PBS again before adding 245 μL of Phalloidin (Ph) (Abcam and Invitrogen, 1:1000) for 30 minutes at RT, protected from light. After 30 minutes, 5 μL of 50X BODIPY stain (Invitrogen) was added for another 25 minutes, making a 1X working concentration of 1:1000 in culture. Finally, 5 μL of 50X 4′,6-diamidino-2-phenylindole (DAPI) (Invitrogen) was added for another 10 minutes, making a 1:1000 working concentration in culture. All stains were diluted in PBS, and extra reagent from the previous stain was used as the basis for the next stain to avoid further dilution. Again, stains were added step-wise to avoid unnecessary disturbances to the cells.

After staining, the cells were rinsed twice with PBS and then kept in 500 μL of PBS.

### BioSpa Cytation 5 Imaging

Wells were imaged at 20X with the BioSpa Cytation 5 Imager with 3 non-overlapping sites imaged per well. The camera model was a Chameleon3 CM3-U3-50S5M. The Gen5 Image Prime 3.12.08 software was used to control imaging parameters and assemble merged images. The channels that were imaged include DAPI (377, 447), TRITC (556, 600), and GFP (469, 525). Exposure and gain parameters were kept consistent in images across the entire experiment.

### Cell Profiler Analysis Workflow

Merged images from the BioSpa Cytation 5 were split into their respective channels using ImageJ2 (v2.14.0/1.54i)^55^. All individual channels were loaded into CellProfiler (v4.2.8)^34^ for Ph and BODIPY analysis. Morphological analysis was done by first identifying all DAPI-positive cells as primary objects using “Minimum Cross–Entropy” as the threshold method, then generating masks of all nuclei. These were the final DAPI objects. Ph-positive regions were measured as secondary objects from the nuclei to create a whole cell mask using the “Watershed – Image” parameter. The size and shape of all Ph objects were measured using the MeasureObjectSizeShape module, which included an area measurement. Area was used to filter out objects that were too small or too large to be considered cells. Filtered Ph objects were saved as a new group of objects and masked. These were the final Ph objects.

BODIPY staining was not measured from DAPI-positive cells, given the placement of the staining with respect to the DAPI-positive nuclei. Instead, BODIPY was measured as a separate primary object using “Minimum Cross – Entropy” as a threshold method. The methods to distinguish and draw dividing lines between clumped objects were intensity and propagate. This ensured that each distinct region of BODIPY positivity was being counted separately, even if they were within the same cell. Similar to Ph objects, BODIPY objects were measured using the MeasureObjectSizeShape module and filtered based on area to remove small puncta. Filtered BODIPY objects were saved as a new group of objects and masked.

Once the filtered object groups for Ph and BODIPY were generated, the MeasureObjectSizeShape function was used again to ensure the Ph measurements and BODIPY areas only include the values from cells that were kept after filtering. Using these revised measurement datasets, the RelateObjects module was used to relate filtered BODIPY objects back to the filtered, final Ph objects. This step is essential because it removes BODIPY-positive regions that do not overlap with Ph. BODIPY-positive, Ph-positive objects were saved as a new dataset and masked. These were the final BODIPY objects. Because CellProfiler now knows to relate BODIPY back to Ph, it can include the Ph measurements next to the respective BODIPY measurements for each cell when the MeasureObjectSizeShape module is used. However, if there were multiple BODIPY-positive areas within a cell, CellProfiler generates the average, rather than the total. To circumvent this and calculate the total area covered by BODIPY, the CalculateMath module was used to multiply the number of BODIPY children per Ph parent by the mean BODIPY area per Ph parent. A separate CalculateMath module was used to calculate the %Area BODIPY by dividing the total BODIPY area by the total Ph parent area per cell and multiplying by 100.

All final DAPI, Ph, and BODIPY objects were converted to images using the ConvertObjectsToImages module. To visualize how much of each Ph object was covered by BODIPY objects, the OverlayObjects function was used with BODIPY objects as the input and Ph as the base objects. To include object numbers in each image, the DisplayDataOnImage module was used. This allowed CellProfiler to add DAPI, Ph, and BODIPY numbers to the respective objects. Additionally, CellProfiler was able to add Ph object numbers to BODIPY images so the BODIPY objects assigned to each Ph parent could be visualized.

Lastly, the ExportToSpreadsheet module was used to save all measurements, and the SaveImages module was used to save all final images. To calculate the percentage of BODIPY-positive cells per image in Excel, the number of BODIPY-positive Ph objects was divided by the total number of Ph objects and multiplied by 100.

All .csv files from CellProfiler were organized in Prism (v10.4.0) and converted from pixels to μm for statistical analysis and graph construction.

All scripts, raw images, and analysis masks can be found on GitHub with the following link: https://github.com/asturno/CellProfiler

## Abbreviations

PBS: Phosphate buffered saline DAPI; 4′,6-diamidino-2-phenylindole
PDL: poly-d-lysine
LPS: lipopolysaccharide
LN2: liquid nitrogen
PFA: paraformaldehyde
FS: fresh
FZ: frozen S1; shake 1
S2: shake 2
Glu: glucose Ctrl; control
Ns: not significant
Ram: ramification
P0-p2: postnatal day 0 - postnatal day 2
IPSC: inducible pluripotent stem cells
FACs: fluorescent activated cell sorting
MAG: magnetic
ATP: adenosine triphosphate
CSF: cerebrospinal fluid
pMG: Primary microglia
LD: Lipid droplet
DMEM: Dulbecco’s Modified Eagle Medium/Nutrient Mixture
FBS: Fetal bovine serum
P/S: Penicillin/streptomycin
DMSO: Dimethyl sulfoxide
Ph: Phalloidin
Max radius: Maximum radius
Ram index: Ramification index
HBSS: Hanks’ Balanced Salt Solution
RT: Room temperature

## Notes

https://github.com/asturno/CellProfiler

### Competing Interest Statement

The authors have declared no competing interest.

### Summary of Updates

The graphical abstract has been uploaded to highlight the main purpose and outcome of this paper.

