## Supplemental Figures for "A validated method for banking mixed glial cultures that yield responsive microglia"

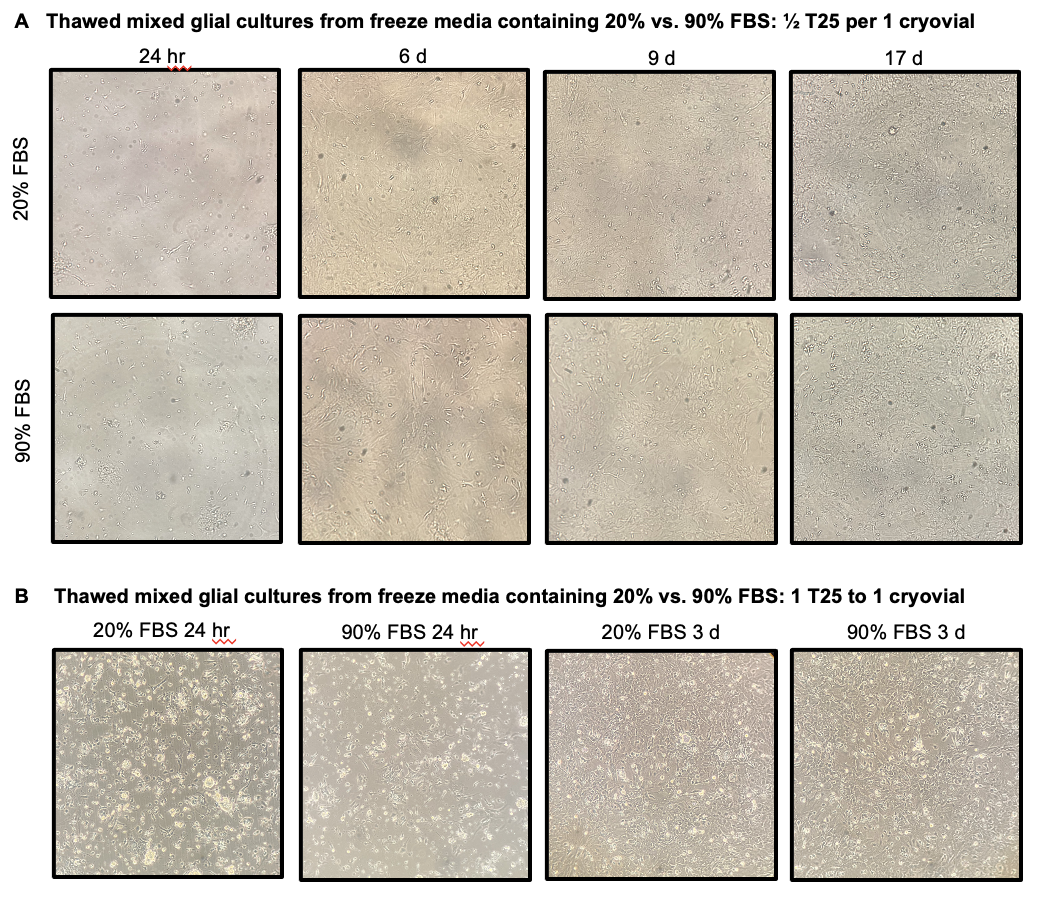

**Figure S1: Mixed glial cultures frozen in media containing 20% and 90% FBS show no differences in viability when thawed and cultured. (A)** Mixed glial cultures frozen in 1 cryovial per ½ T25 in media containing 20% or 90% FBS 24 hr, 6 d, 9d, and 17d after thaw. **(B)** Mixed glial cultures frozen in 1 cryovial per 1 T25 containing 20% or 90% FBS 24 hr and 3d after thaw.

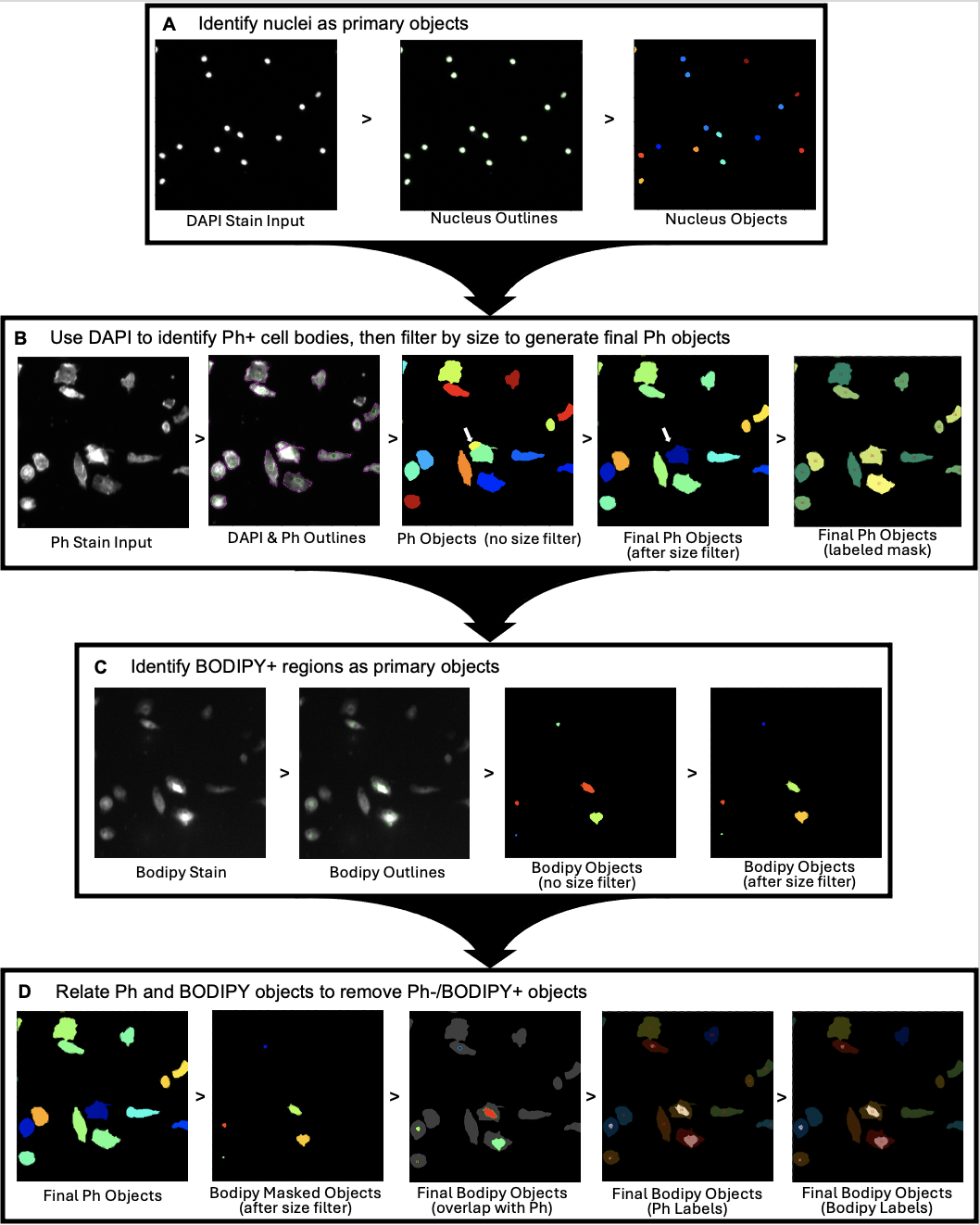
**Figure S2: Cell Profiler workflow for identification of Ph and BODIPY-positive regions in pMG. (A)** Identification of live nuclei from DAPI+ regions. **(B)** Masking and filtering of Ph+ objects propagated from DAPI+ nuclei for pMG size and shape analysis. **(C)** Identification of BODIPY+ regions to extract areas of neutral lipid accumulation as a proxy for LDs. Although BODIPY masks were filtered for unusually small or large positive regions, their occurrences were rare and are not pictured here. **(D)** Use of final Ph objects and filtered BODIPY objects to generate final BODIPY objects and ensure only Ph+ and BODIPY+ objects are used for LD accumulation analysis.

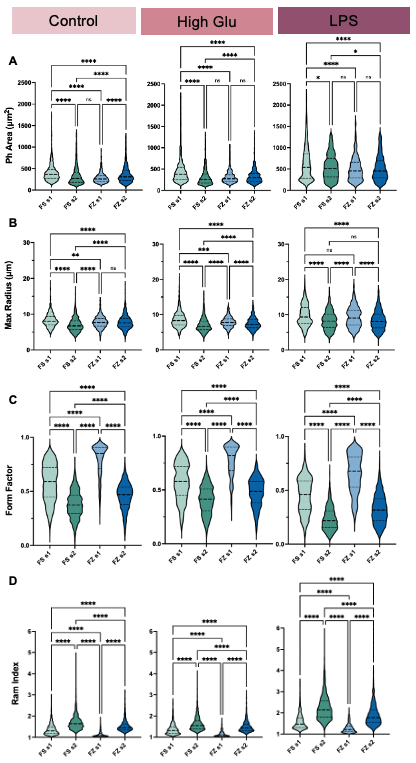

**Figure S3: Absolute responses of pMG morphology to control, High Glu, and LPS conditions differ across FS-FZ status and shake number.** Microglia cultured from FS s1, FS s2, FZ s1, FZ s2 mixed glial cultures exhibit differences in their absolute measurements of **(A)** Ph area (µm^2^), **(B)** max radius (µm), **(C)** form factor, and **(D)** ram index. Data are presented as quartiles (n = 300-1,500 cells per group), *P < 0.05, **P < 0.01, ***P < 0.001, ****P < 0.0001 (Kruskal-Wallis nonparametric test with Dunn's multiple comparisons test), ns = not significant.

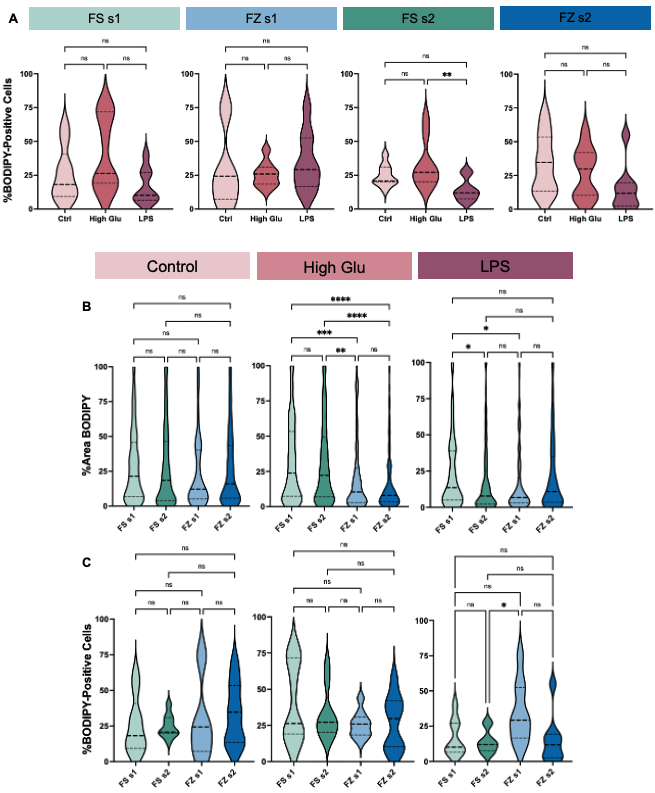

**Figure S4: pMG have variable responses in the %BODIPY+ cells and %Area BODIPY across FS-FZ status and shake number in response to High Glu and LPS.** **(A)** pMG exhibit no significant changes in %BODIPY+ cells in response to control, High Glu, and LPS conditions. The absolute responses of pMG cultured from FS s1, FS s2, FZ s1, FZ s2 mixed glial cultures exhibit differences in their absolute measurements of **(B)** %Area BODIPY and **(C)** %BODIPY+ cells. Data are presented as quartiles (n = 8-9 images per group), *P < 0.05, **P < 0.01, ***P < 0.001, ****P < 0.0001 (Kruskal-Wallis nonparametric test with Dunn's multiple comparisons test), ns = not significant.
